# The neural response to the temporal fine structure of continuous musical pieces is not affected by selective attention

**DOI:** 10.1101/2021.01.27.428483

**Authors:** Octave Etard, Rémy Ben Messaoud, Gabriel Gaugain, Tobias Reichenbach

**Affiliations:** Department of Bioengineering and Centre for Neurotechnology, Imperial College London, South Kensington Campus, SW7 2AZ, London, UK

## Abstract

Speech and music are spectro-temporally complex acoustic signals that a highly relevant for humans. Both contain a temporal fine structure that is encoded in the neural responses of subcortical and cortical processing centres. The subcortical response to the temporal fine structure of speech has recently been shown to be modulated by selective attention to one of two competing voices. Music similarly often consists of several simultaneous melodic lines, and a listener can selectively attend to a particular one at a time. However, the neural mechanisms that enable such selective attention remain largely enigmatic, not least since most investigations to date have focussed on short and simplified musical stimuli. Here we study the neural encoding of classical musical pieces in human volunteers, using scalp electroencephalography (EEG) recordings. We presented volunteers with continuous musical pieces composed of one or two instruments. In the latter case, the participants were asked to selectively attend to one of the two competing instruments and to perform a vibrato identification task. We used linear encoding and decoding models to relate the recorded EEG activity to the stimulus waveform. We show that we can measure neural responses to the temporal fine structure of melodic lines played by one single instrument, at the population level as well as for most individual subjects. The neural response peaks at a latency of 7.6 ms and is not measurable past 15 ms. When analysing the neural responses elicited by competing instruments, we find no evidence of attentional modulation. Our results show that, much like speech, the temporal fine structure of music is tracked by neural activity. In contrast to speech, however, this response appears unaffected by selective attention in the context of our experiment.

## Introduction

Music is a fascinatingly complex acoustic stimulus. Listeners can follow multiple melodic lines played by different instruments by separating them on the basis of characteristics such as pitch and timbre (Cross et al., 2008). However, the neural mechanisms that group the sounds in music into distinct melodic lines, forming distinct auditory streams, and allow attention to be directed to one of the lines remain largely unknown (Albert S Bregman, 1994). This is partly due to the difficulty in assessing the neural processing of real-world acoustic signals that have a much richer structure than the simple pure tones and short simplified music patterns that have traditionally dominated research in auditory neuroscience.

A better understanding of the neural mechanisms of music processing may emerge from combining statistical models with neuroimaging. Recent studies have indeed shown how these methods can relate key features of a complex sound such as speech to electrophysiological recordings and inform on the neural mechanisms of speech processing (Di Liberto et al., 2015; Ding & Simon, 2012a, 2014; Wöstmann et al., 2017). For example, cortical activity has been found to track slow (< 8Hz) amplitude fluctuations in speech (Ding & Simon, 2012b; Edmund C. Lalor & Foxe, 2010; Nourski et al., 2009; Pasley et al., 2012), while subcortical as well as, presumably to a lesser degree, cortical responses emerge to the higher frequency (> 80 Hz) stimulus structure (Bidelman, 2018; Coffey et al., 2016; Etard et al., 2019; Forte et al., 2017; Maddox & Lee, 2018). The temporal fine structure of speech originates from the periodic opening and closing of the vocal folds at the so-called fundamental frequency. The spectrum of these voiced speech parts is therefore dominated by the fundamental frequency as well as its many higher harmonics, leading to a pitch perception in the listeners.

Understanding how the brain can focus on a single instrument amongst others relates to a major challenge in auditory neuroscience, the cocktail party problem. This problem acquired its name from the observation that humans do remarkably well at understanding a target speaker in a noisy environment such as in a busy restaurant or in a loud bar (Cherry, 1953; Haykin & Chen, 2005). A recent study showed that neural responses to the pitch of continuous speech are stronger when the stimulus is attended rather than ignored (Forte et al., 2017). This result suggests that the pitch of a speaker could be used by the brain to perceptually segregate the speech signal from background noise, a finding that agrees with previous psychophysical studies that have found it easier to differentiate two concurrent speech signals if their fundamental frequencies differ (de Cheveigné et al., 1997; Madsen et al., 2017).

Musical tones are similarly characterized by a fundamental frequency and higher harmonics, resulting in a characteristic temporal structure that causes a pitch perception. The proximity of fundamental frequencies of subsequent tones has been found to aid the formation of an auditory stream (A. S. Bregman et al., 1990; Oxenham, 2008). Consequently, just as the neural tracking of temporal fine structure could help listeners attend to a voice in background noise, such a neural mechanism may aid with attending to a particular melodic line (Micheyl & Oxenham, 2010).

Here we investigated this hypothesis by using linear models to assess neural responses to the temporal fine structure of continuous melodic lines. We first presented volunteers with continuous classical Bach pieces while recording their brain activity through a bipolar EEG montage. We then related the neural activity to the stimulus waveforms using encoding and decoding methods. To assess a putative effect of selective attention on these neural activities, we also presented the volunteers with two competing instruments, a guitar and a piano, that were playing two different melodic lines simultaneously. Subjects were asked to selectively attend to one of the two lines, and we contrasted the neural responses to each instrument when it was attended to when it was ignored.

## Methods

Due to multiple nonlinearities in the auditory periphery, both the temporal fine structure and the envelope of the stimuli are represented in neural responses. This encoding has traditionally been investigated in humans by studying time-locked responses to transient or periodic features of repeated short sound tokens such as clicks, pure or complex tones, syllables and words (Skoe & Kraus, 2010). These paradigms typically present a particular stimulus as well as its opposite waveform many times. The neural responses to each polarity are then summed to emphasise responses to the envelope or subtracted to emphasise response to the temporal fine structure (Aiken & Picton, 2008; Krizman & Kraus, 2019). Here we used continuous, long stimuli to derive auditory neural responses to their temporal fine structure using linear convolutive models.

### Experimental design and statistical analysis

Seven of Bach’s Two-Part Inventions were used in this study. Each Two-Part Invention is a short keyboard composition that consists of two melodic lines: one played by the left hand, and one by the right. We synthesized the stimuli in GarageBand (Apple, U.S.A) from Musical Instrument Digital Interface (MIDI) files, with the left hand being played by a piano and the right by a guitar. To assess the attention of subjects to a particular melodic line, vibratos were inserted in both lines.

Volunteers were presented with two type of stimuli. The first type, “Single Instrument” (SI), consisted of one single instrument, piano or guitar, that played one melodic line. The second type “Competing Instruments” (CI), contained both melodic lines of a Two-Part Invention, one played by the piano and the other by the guitar.

The different stimuli were presented in blocks (figure 1). Each block contained one SI stimulus and one subsequent CI stimulus, both of which were obtained from the same Two-Part Invention. During the CI stimulus, the volunteers were asked to selectively listen to the instrument that they heard before in the SI stimulus. They were also asked to identify the vibratos embedded into that melodic line.

**Figure 1:**
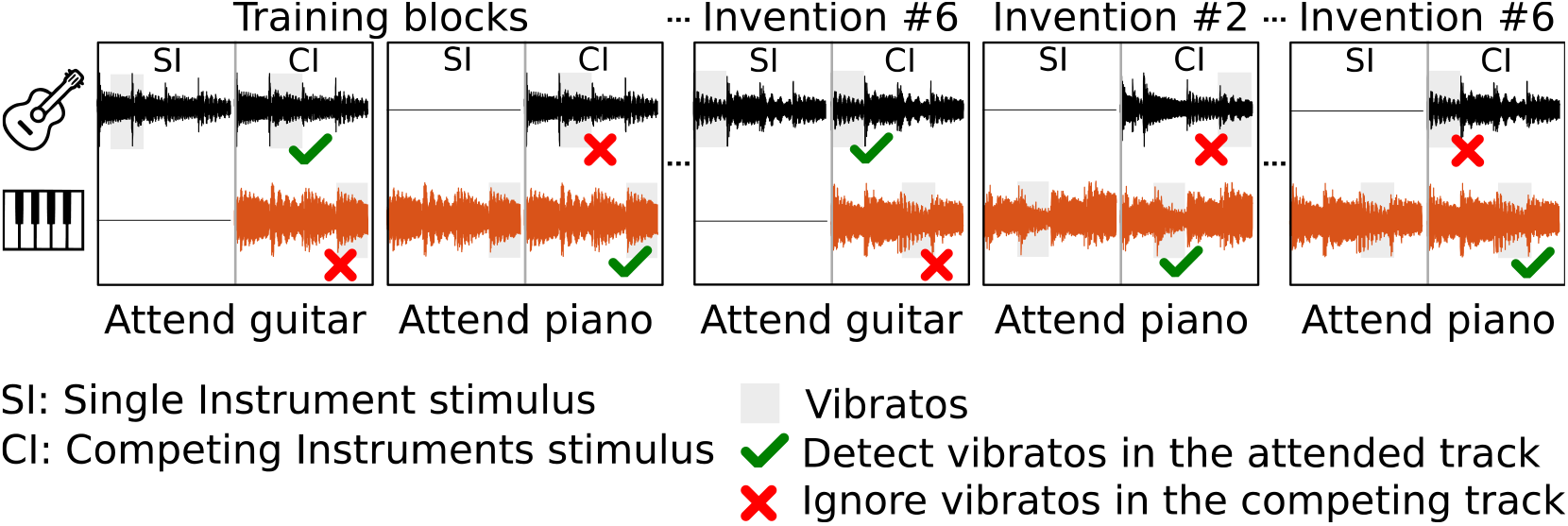
Schematic representation of the experiment. Volunteers were presented with continuous classical music pieces, Bach’s Two-Part Inventions, that consisted of either a single melodic line (SI) or of two melodic lines (CI). Each melodic line was played either by a guitar or by a piano. In the CI stimuli, each melodic line was played by a different instrument. Vibratos were inserted into the acoustic waveforms of each melody (grey shading). In the CI condition, the subjects had to attend to one of the two instruments and identify the corresponding vibratos (green tick marks) while ignoring the other instrument and its vibratos (red crosses). The stimuli were presented in blocks composed of a SI stimulus followed by a CI stimulus during which the subject was asked to attend to the instrument that they heard before in the SI stimulus. The attended instrument was alternated between blocks, and each block was played twice such that the attended instrument differed in the two presentations. The volunteers’ neural responses were recorded throughout the experiment through bipolar two-channel EEG recordings.

Blocks with the SI stimulus played by the piano alternated with those played by the guitar. Each of the seven Two-Part inventions was presented twice: once with the SI stimulus played by the guitar, and once with the SI stimulus played by the piano. Each participant therefore heard seven SI stimuli played by the guitar, seven SI stimuli played by the piano, and fourteen CI stimuli.

All participants were initially presented with the same two training blocks, one with a SI stimulus played by the guitar and one played by the piano, that corresponded to the same invention. These stimuli presentations served to familiarise the subjects with the task of attending to one melodic line in the CI stimulus and to identify the embedded vibratos. These training blocks were excluded from further analysis, leaving six inventions in each condition.

The presentation order of the remaining blocks was pseudo-randomised across participants. In the second presentation of a given CI stimulus, a subject was asked to attend to the instrument they ignored in the first presentation. Two consecutive blocks did not correspond to the same invention. Each participant therefore heard each CI stimulus twice, but attended a different instrument in each presentation. Whether a participant was initially asked to attend to the guitar or the piano was randomly decided. The participant’s neural responses were measured through scalp electroencephalography (EEG) with a two-channel bipolar montage (head vertex minus mastoids).

We used encoding and decoding approaches (linear forward and backward models) to relate the acoustic stimuli to the recorded neural data. We specifically investigated the neural representation of the temporal fine structure by using the stimulus waveform as a feature. We first established that we could indeed record a significant neural response to this feature, by comparing the neural responses to a null distribution at the level of individual subjects as well as on the population level. We then studied the time course of the response in the region between 0 to 45 ms, using both forward and backward models. Finally, we investigated a putative attentional modulation of this neural response through contrasting the encoding of each instrument in the neural data when attended versus ignored. We used conservative filters to reduce distortions to the neural responses and their latencies, but verified that our results, and in particular the ones related to attention, did not change with stronger filtering (data not shown).

### Code and data availability

The analysis presented in this manuscript was implemented using MATLAB (R2019b, The MathWorks Inc.) with the EEGLAB toolbox (Delorme & Makeig, 2004). The linear forward and backward models were trained using the LMpackage (github.com/octaveEtard/LMpackage). The raw data as well as analysis code and processed data required to reproduce the results presented here were made available (https://github.com/octaveEtard/EEGmusic2020; https://zenodo.org/record/4470135).

### Participants

17 volunteers (aged 23.8 ± 2.9 year, 9 females) participated in this experiment. The number of participants was chosen based on previous studies investigating similar neural responses to continuous speech (Etard et al., 2019; Forte et al., 2017). All participants were right-handed, had no history of auditory or neurological impairments, and provided written informed consent. The experimental procedures were approved by the Imperial College Research Ethics Committee.

### Music stimuli

To generate neural responses to each instrument that were of similar magnitude, the notes of the guitar were lowered by one octave so that their fundamental frequencies fell below 500 Hz. They remained nonetheless somewhat higher than those of the piano notes (figure 2, A, B). The music stimuli were synthesized from MIDI files to generate wav files. These were then processed using MATLAB to apply vibratos to ten segments in each melodic line. Each vibrato was constructed by introducing a sinusoidal warp at a modulation frequency of *f*_*m*_= 8 Hz on the waveform of a single note. The onset and offset times of the notes were obtained from the MIDI files using the Miditoolbox for Matlab (Eerola & Toiviainen, 2004). The notes were selected such that the onsets of any two vibratos in a given piece, whether both played by the same or different instruments, were separated by at least one second.

**Figure 2:**
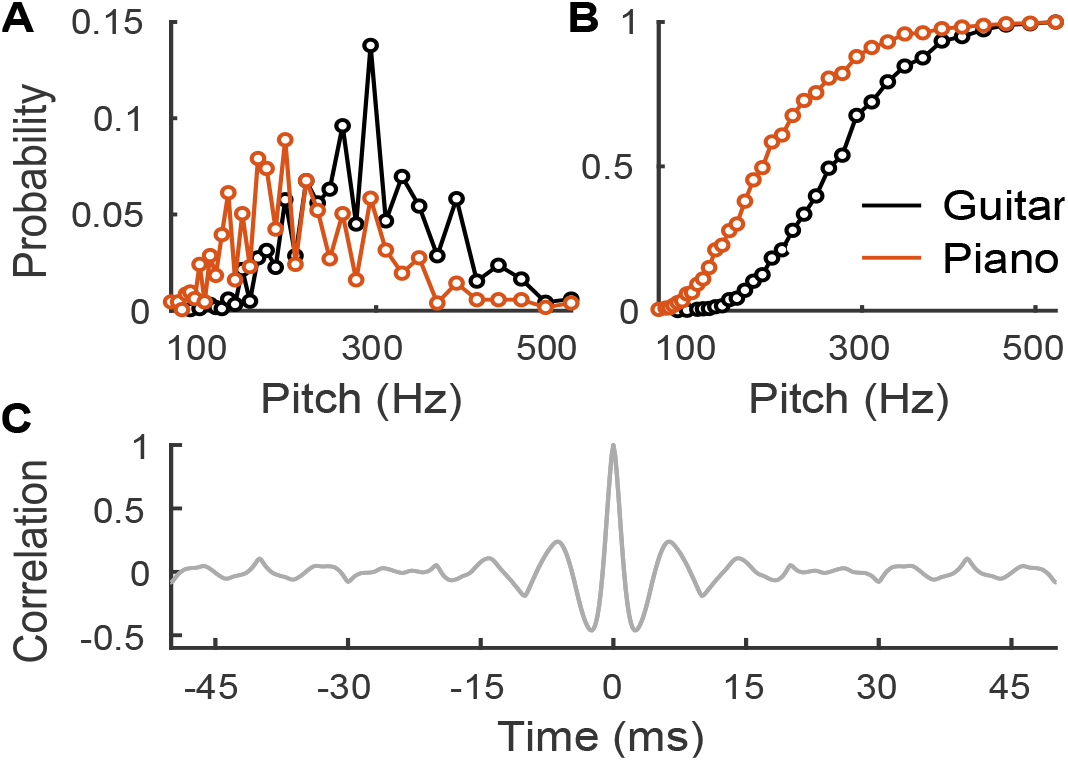
Properties of the acoustic stimuli and of the filtered EEG data. (**A**), The probability mass function of the fundamental frequency of the notes peaked at around 196 Hz for the piano (red), and at about 294 Hz for the guitar (black). (**B**), The cumulative distribution of the fundamental frequency of the notes showed likewise that most fundamental frequencies lied between 100 Hz and 400 Hz, with the distribution of the guitar notes being shifted to somewhat higher frequencies. (**C**), To eliminate frequencies below the range of the fundamental frequencies, the EEG data was high-pass filtered above 130 Hz. The filtered EEG data consequently displayed some periodicity and correlation in time as evident from its auto-correlation function.

The waveforms of the CI stimuli, *w*_*mixed*_, were constructing by normalising and mixing the waveform *w*_*g*_ of the guitar and the waveform *w*_*p*_ of the piano according to their root-mean-square values (RMS):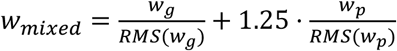. The mixing parameter of 1.25 for the piano was chosen following a small pilot study to balance the difficulty in attending either the guitar or the piano.

The duration of the seven Two-Part Inventions was, taken together, 11.2 minutes. In the SI conditions, only the first half of the corresponding invention was played.

### Behavioural task

In the CI condition, the subjects were instructed to attentively listen to one instrument while ignoring the other. They were also asked to classify the vibratos they heard by pressing a key to indicate the ones that belonged to the attended instrument. A key press within two seconds after the onset of a vibrato in the attended or ignored instrument was classified respectively as “true positive” (TP) or “false positive” (FP). Key presses outside of these ranges were classified as “unprompted” and were not analysed further. Due to a technical error, behavioural data was not recorded for one subject, and only the results for the 16 remaining subjects were analysed. The sensitivity index d-prime was computed for each subject when attending to the guitar and the piano, and it was compared between the two conditions at the population level using a two-tailed paired Wilcoxon signed rank test. Moreover, for each condition the TP rate (TPR) was compared to the FP rate (FPR), and the TPR and FPR were compared between conditions at the population level using two-tailed paired Wilcoxon signed rank tests with FDR correction for multiple comparisons (four tests).

### Neural data acquisition and stimulus presentation

Scalp EEG was recorded through five passive Ag/AgCl electrodes (Multitrode, BrainProducts, Germany). Two electrodes were positioned on the cranial vertex (Cz), and two electrodes were placed on the left and right mastoid processes. A ground electrode was placed on the forehead. The impedance between each electrode and the skin was reduced below 5 kOhm using abrasive electrolyte gel (Abralyt HiCl, Easycap, Germany). One vertex electrode was paired with the left mastoid electrode, and they were connected to, respectively, the non-inverting and inverting ports of a bipolar amplifier (EP-PreAmp, BrainProducts, Germany). The remaining vertex and mastoid electrodes were similarly connected to a second identical amplifier. The output of each bipolar pre-amplifier was fed into an amplifier (actiCHamp, BrainProducts, Germany) and digitized with a sampling frequency of 5 kHz, thus yielding two electrophysiological data channels. The audio stimuli were simultaneously recorded at 5 kHz by the amplifier through an acoustic adapter (Acoustical Stimulator Adapter and StimTrak, BrainProducts, Germany). This channel and independent analogue triggers delivered through an LPT port were used to temporally align the EEG data and stimuli through cross-correlation. The stimuli were delivered diotically at a comfortable loudness level through insert tube earphones (ER-3C, Etymotic, USA) to minimise stimulation artifacts. These earphones introduced a 1 ms delay that was compensated for by shifting the neural data forward in time by 1ms.

### EEG data filtering

To analyse the neural responses to the temporal fine structure of the stimuli, the EEG data was high-pass filtered above 130 Hz (windowed-sinc filters, Kaiser window, one pass forward and compensated for delay; cut-off: 115 Hz, transition bandwidth: 30 Hz, order: 536). These filters rejected lower-frequency neural activity but reduced the temporal precision of the data, as evidenced by the auto-correlation function of the filtered EEG data (figure 2C). Notably, they were non-causal filters that spread responses in both temporal directions.

### Stimulus representation

Since the vibratos might lead to neural responses deviating from the ones elicited by the rest of the tracks, the parts of the stimulus waveforms that corresponded to them were replaced with zeros to create the stimulus representations (features) used in the encoding and decoding models. These waveforms were then low pass filtered and resampled from 44.1 kHz to 5 kHz, the sampling frequency of the EEG data, using a linear phase FIR anti-aliasing filter (windowed-sinc filter, Kaiser window, one pass forward and compensated for delay; cut-off: 2,250 Hz, transition bandwidth: 500 Hz, order: 14,126).

### Encoding models

We used regularised linear forward models to derive the neural response to the stimulus waveform. In these convolutive encoding models, the measured EEG response *e* is modelled as *e*(*t*) = (*r* * *s*)(*t*) + *n*(*t*), where *s* is the stimulus waveform, *r* the neural response or Temporal Response Function (TRF), *n* is noise, and * is the convolution symbol. In practice, assuming a non-zero response in a time interval (*τ*_*min*_; *τ*_*max*_) only, and with discrete data, the EEG activity *e*_*i*_(*t*_*n*_) at channel *i* ∈ {1; 2} and at time *t*_*n*_ can be estimated as 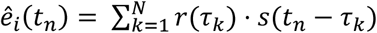, with *τ*_1_ = *τ*_*min*_ and *τ*_*N*_ = *τ*_*max*_. Given the bipolar montage we used, as well as the diotic stimulus presentation, we did not expect any difference between the two EEG channels and assumed the same neural response for both.

The model was estimated for time lags spanning *τ*_*min*_ = −100 ms to *τ*_*max*_ = 45 ms. A population-averaged TRF *r* was fitted using ridge regression coupled with a leave-one-subject-out and leave-one-data-part out cross-validation (Crosse et al., 2016; Hastie et al., 2009; E. C. Lalor et al., 2009). The model was fitted using the data corresponding to all the stimulus parts bar one and all the subjects bar one, and evaluated on the left-out data part for the left-out subject. The stimulus part and the subject used for testing were hence not seen by the model during training. This constituted one cross-validation fold. The left-out subject and the left-out data parts were then iterated until all combinations were exhausted, for a total of 17 × 6 = 102 folds. The validation performance of the model was quantified by dividing the predicted neural response 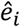 and the measured EEG activity *e*_*i*_ from the testing data in each fold into 10-s long segments, and by computing Pearson’s correlation coefficients between each segment. The correlation coefficients thus obtained were then averaged over all cross-validation folds as well as over all EEG channels.

The performance was assessed for models corresponding to 25 normalised regularisation coefficients *λ*_*n*_ that were distributed uniformly on a logarithmic scale between 10^−6^ and 10^6^. The regularisation coefficient was thereby *λ* = *λ*_*n*_ · *m*, with *m* the mean eigenvalue of the predictor’s auto-correlation matrix (Biesmans et al., 2017). The model yielding the highest reconstruction performance was chosen as representing the neural response. To assess the significance of the obtained TRFs, the negative, non-causal part of the response, −100 ms to 0 ms, was used to construct a null distribution. For each instrument, a Gaussian distribution was fitted to the pooled data points from the negative part of the response. From the distribution we determined the *p*-values of all the points in the positive part of the response (0 ms to 45 ms), and applied an FDR correction for multiple comparison over time points and instruments.

To ascertain the relative contributions of the onset and of the sustained parts of the notes to the neural response, we created a new representation of the stimuli in which the note onsets were suppressed. This was achieved by multiplying the original stimulus waveforms by a 60-ms window *w* centred on each note onsets, with *w*(*t*) = 1 − *h*(*t*) and *h* representing a 60-ms Hann window. Forward models were then derived for the original stimuli and their onset-suppressed versions for the two SI conditions taken together, by pooling the data from both instruments. These two models were fitted and their significance was ascertained as described above, that is, by comparing the causal part to the null models, with FDR correction for multiple comparison over time points and over the two models. In the cross-validation procedure, two data parts, one from each SI condition and corresponding to the same invention, were left out at each stage.

### Decoding models

We also used backward models to reconstruct the stimulus waveform *s* as a linear combination of the neural activity *e*_*i*_ on each channel *i* at different time lags: 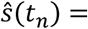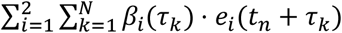, with *τ*_*min*_ ≤ *τ*_*k*_ ≤ *τ*_*max*_. The coefficients *β* were trained for each subject independently, using ridge regression with a leave-one-part-out cross-validation and a normalised regularisation coefficient *λ*_*n*_ = 10^−0.5^ (Biesmans et al., 2017). As with the forward models, the performances of the backward models were measured through computing the correlation coefficients between the reconstructed stimulus and the actual one on 10-s long segments of the testing data. The set of correlation coefficients pooled from all cross-validation folds for a given participant was used when performing statistical testing at the level of individual subjects, and the corresponding average correlation coefficient was used for each subject when testing at the population level. The performances of the models were thereafter used to quantify the neural encoding of each stimulus for a given reconstruction time window *τ*_*min*_ − *τ*_*max*_.

### Significance of the stimulus reconstruction

The neural encoding of the SI stimuli for each instrument was measured through the backward models using reconstruction time windows of equal duration but centred on different delays. To establish the significance of the stimulus reconstruction procedure at the level of individual subjects, a window of delays between *τ*_*min*_ = −15 ms and *τ*_*max*_ = 0 ms was used to provide a null distribution for each subject. The neural encoding in the window of interest, from *τ*_*min*_ = 0 ms to *τ*_*max*_ = 15 ms, was compared to the null distribution for each subject using one-tailed paired Wilcoxon signed rank tests with FDR correction for multiple comparisons over subjects and instruments. Significance was also derived at the population level using the mean correlation coefficients for each subject from the null window of negative delays to create a null population-level distribution. To test the time windows in which a significant response could be detected, the mean reconstruction accuracies from three windows of interest (0 to 15 ms; 15 to 30 ms; 30 to 45 ms) were compared to this null distribution using one-tailed paired Wilcoxon signed rank tests with FDR correction for multiple comparisons over windows and instruments.

Since the guitar and piano waveforms formed pairs derived from the same inventions, and although their frequency contents were different, one may wonder whether one instrument could be predicted from the other, and in turn whether the neural responses to one instrument could be predicted or used to decode the other one. To address this question, we trained linear backward models that sought to reconstruct the waveform of one instrument from the neural data that was recorded when the other instrument from the same invention was played in the SI conditions (0 to 15 ms reconstruction window). The model performance was then compared to the null distribution previously described (obtained from a −15 to 0 ms reconstruction window) at the population level, using one-tailed paired Wilcoxon signed rank tests.

### Competing conditions, attended and ignored instruments

In the CI conditions, we trained backward models to reconstruct the waveform of either the attended or the ignored instrument independently, using a window of temporal delays from *τ*_*min*_ = 0 ms to *τ*_*max*_ = 15 ms as detailed above. We then compared the neural encoding of each instrument, when attended and when ignored, at the population level, using two-tailed paired Wilcoxon signed rank tests with FDR correction for multiple comparisons over instruments.

We also used forward models reconstructing the neural activity as the sum of two neural responses, one to the attended instrument and one to the ignored one. In this instance, the EEG response *e* is modelled as *e*(*t*) = (*r*_*A*_ * *s*_*A*_)(*t*) + (*r*_*I*_ * *s*_*I*_)(*t*) + *n*(*t*), where *s*_*A*_ and *s*_*I*_ are the attended and ignored stimulus waveforms, and *r*_*A*_ and *r*_*I*_ the corresponding TRFs. In a similar manner to the procedures previously described, population-averaged TRFs were fitted using ridge regression coupled with a leave-one-subject-out and leave-one-data-part out cross-validation for time lags spanning *τ*_*min*_ = −100 ms to *τ*_*max*_ = 45 ms on the pooled data from the two CI conditions. To assess the presence of a putative attentional modulation in the obtained TRFs, the distribution of amplitude across subjects was compared between the attended and ignored TRFs for each time point in the 0 ms to15 ms region of interest (two-tailed paired Wilcoxon signed rank tests with FDR correction for multiple comparisons over time points).

## Results

We asked volunteers to attend to continuous musical pieces consisting of either one single instrument (SI) or of two competing instruments (CI) while we recorded their neural activity using EEG (figure 1). We first sought to analyse the neural response to the temporal fine structure of a single melodic line. To this end, we computed a linear forward model to derive neural responses to the stimulus waveform at the population level in the SI conditions (figure 3A). The temporal response functions that we obtained for the two instruments were qualitatively similar to each other. They displayed a major significant response at a latency of 7.6 ms, as well as a minor positive peak at 2.2 ms, with sidelobes reminiscent of the EEG auto-correlation function (figure 2C).

**Figure 3:**
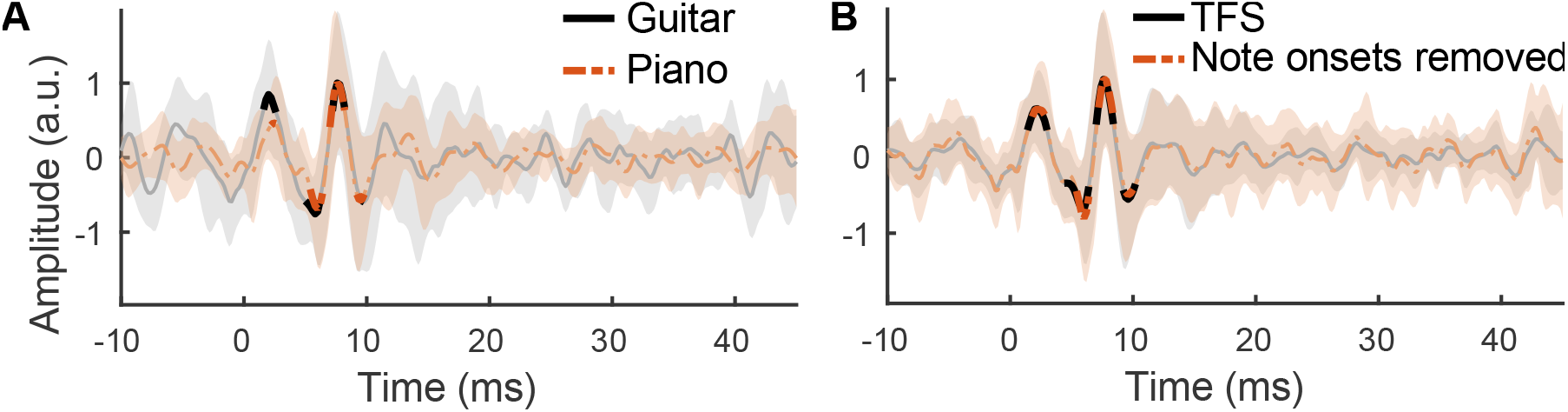
Temporal Response Functions (TRFs). (**A**), We obtained TRFs on the population level from forward models that predicted the neural responses from the stimulus temporal fine structure in the SI conditions for the guitar (black) and for the piano (red). Shaded regions denote plus/minus one standard deviation across subjects around the mean TRFs. Significant regions (thick lines) emerged at similar latencies for the guitar and piano, with a first peak at 2.2 ms, followed by a main positive peak at 7.6 ms. (**B**), We also computed TRFs for both instruments taken together from stimulus waveforms in which the note onsets where removed (red). The obtained TRFs exhibited nonetheless the same significant peaks as the TRFs from the original temporal fine structure feature (TFS, black), indicating that the neural response was not influenced by the note onsets.

The neural response to temporal fine structure may be related to the well-established frequency-following response (FFR). Because the latter is known to first exhibit a response to a stimulus onset and to then follow the sustained features, we explored the relative contributions of the note onsets and their sustained oscillations to the neural response. We therefore trained a forward model with stimulus waveforms in which the note onsets were suppressed (figure 3B). The obtained temporal response functions had similar significant regions, and resembled the temporal response functions to the original stimulus waveforms. Moreover, the causal parts of the two temporal response functions, those with positive delays, were highly correlated (*r* = 0.96).

As an alternative method to the forward models, we then also used decoding models that reconstructed the stimulus waveforms based on the EEG data. We computed these models for each subject in the SI condition. To ascertain the statistical significance of the reconstructions, we used a window from −15 ms to 0 ms to provide a null distribution of performance. Compared to this chance level, we found that a significant reconstruction accuracy could be obtained for most subjects when using time lags from 0 to 15 ms for both guitar and piano (figure 4A). Indeed, significant reconstructions of the guitar waveforms were obtained in 11 out of 17 subjects (*p* ≤ 0.05), in 10 subjects for the piano waveforms, and in 8 subjects for both types of stimuli. The reconstructions of the waveforms for the guitar and for the piano were also significant at the population level (figure 4B; guitar: *p* = 1.6 · 10^−2^; piano: *p* = 1.54 · 10^−3^). Finally, on the population level, when assessing the statistical significance of the stimulus reconstructions using each of three windows of interest (0 to 15 ms, 15 to 30 ms and 30 to 45 ms), we found that only the window from 0 to 15 ms yielded a significant reconstruction accuracy, for either instrument (figure 4C).

**Figure 4:**
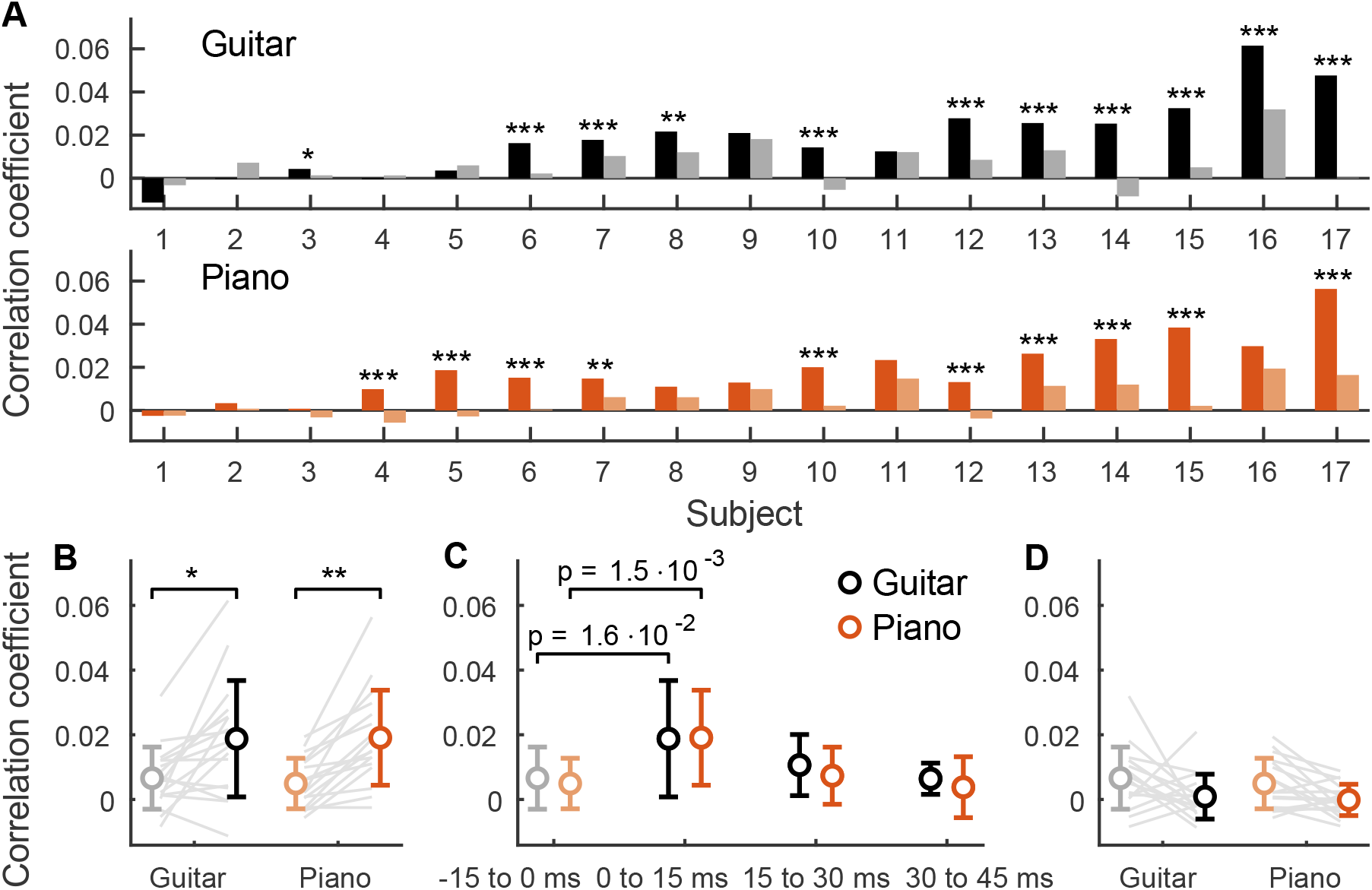
Backward models that reconstruct the stimulus waveform from the EEG data in the SI condition. (**A**), In most subjects, the backward models gave a stimulus reconstruction that had a significantly larger correlation (dark colour) with the original waveform than a null model (light colour). The volunteers were sorted by mean performance, and asterisks indicate *p*-values (*: *p* ≤ 0.05, **: *p* ≤ 0.01, ***: *p* ≤ 0.001). (**B**), The mean reconstruction accuracy for each subject was used to test the significance of the reconstruction at the population level. Both the guitar and the piano stimuli could be reconstructed significantly better from the EEG recordings than from null models. (**C**), We also assessed the reconstruction of the backward models using three windows of temporal delays: 0 ms to 15 ms, 15 ms to 30 ms, and 30 ms to 45 ms (dark colours), and compared them to a null model obtained from the negative delays of −15 ms to 0 ms (light colours). Only the temporal window of 0 ms to 15 ms allowed for a stimulus reconstruction that was significantly better than that of the null model. (**D**), Reconstructing one instrument waveform using the EEG recorded during the presentation of the other instrument (0 to 15 ms window; dark colours) did not yield significant performances as compared to the null model derived by using negative delays (−15 to 0 ms; light colours).While the two instrument waveforms formed pairs corresponding to an invention, the waveform of one instrument could not be predicted from the neural responses to the other instrument.

As the stimuli we used were derived from the left and right hands of inventions, one may wonder whether two instrument waveforms derived from the same piece are independent, and whether the neural responses to one instrument could be used to decode the other one. This is particularly relevant in the context of the attention experiment where such an effect could obscure a putative attentional modulation. However, the stimulus reconstruction accuracy when mismatching the EEG – stimuli pairs in such a way (0 to 15 ms reconstruction window) was not significant as compared to the null distribution using matched EEG – stimuli pairs and a −15 to 0 ms reconstruction window (figure 4D; guitar: *p* = 0.99; piano: *p* = 0.96).

Armed with the ability to measure neural responses to the temporal fine structure of the notes in a particular melody, we then investigated whether this response was affected by selective attention. To this end, we analysed the CI stimuli, in which the participants had to attend selectively to one instrument while ignoring the other. We monitored attention by asking the volunteers to classify vibratos inserted into the melodic line played by the target instrument. The participants exhibited varied performances on this task, however they all had an average performance that was better than that of a random observer, as shown by their receiver operating characteristics, when selectively attending to either of the two instruments (figure 5A). Accordingly, at the population level, the TPR was significantly larger than the FPR when attending to either instrument (*p* < 10^−3^ for guitar and piano). The sensitivity index d’ was significantly larger when attending to the piano than when attending to the guitar (*p* = 1.5 · 10^−2^), with an average value 2.0 and 1.5, respectively (figure 5B). The TPR did not differ significatively between the two CI conditions (*p* = 0.88; figure 5C), but the FPR was significantly higher when the subjects were attending to the guitar compared to the piano (FPR: *p* = 1.79 · 10^−3^, figure 5D).

**Figure 5:**
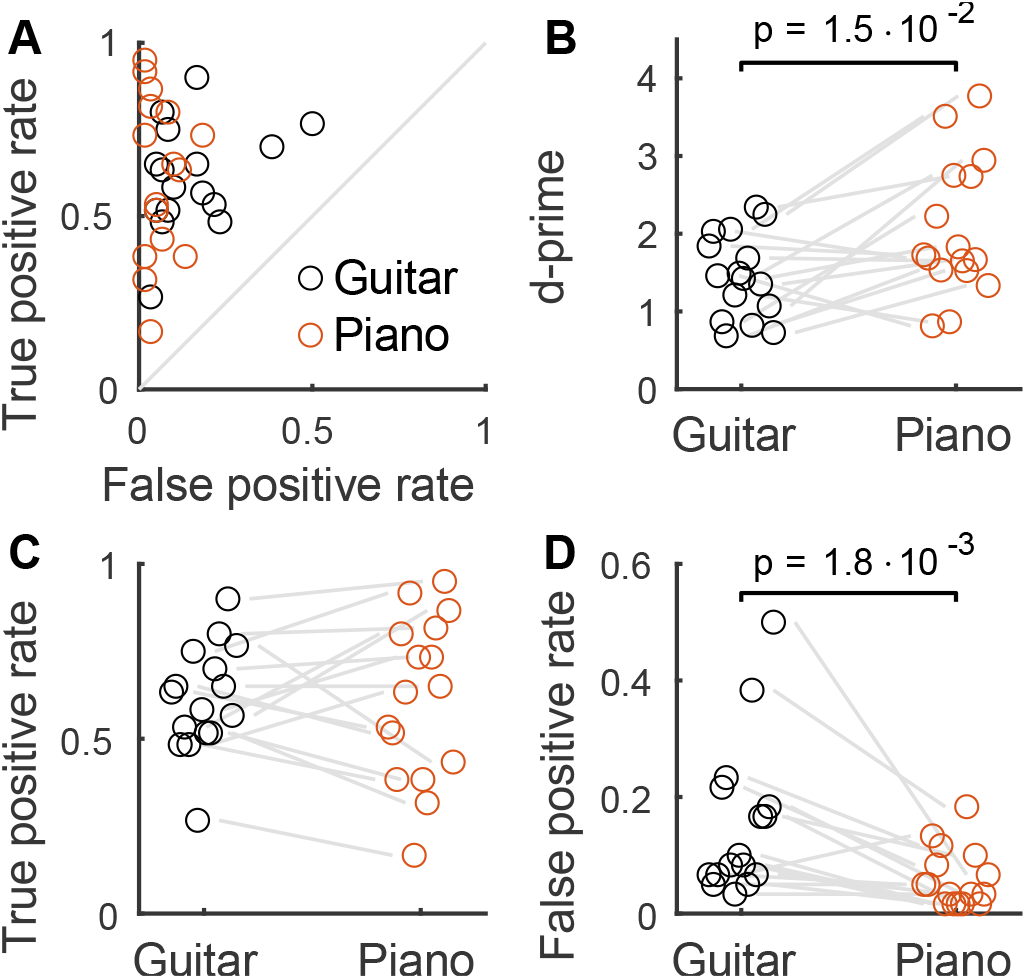
Behavioural results for the vibrato classification, task. Each circle represents a subject. (**A**), The receiver operating characteristics (ROC) shows that each subject performed above chance level in the CI condition, both when attending to the guitar (black) and when attending to the piano (red). (**B**) The average sensitivity index d’ was significantly larger when attending to the piano than to the guitar (*p* = 1.5 · 10^−2^) with an average value of 2.0 and 1.5, respectively. (**C**), The rate of true positives was similar when attending to the guitar and then attending to the piano. (**D**), Attending to the guitar led to more false positives than attending to the piano (*p* = 1.8 · 10^−3^).

In order to test for a putative attentional modulation of the encoding of the stimulus temporal fine structure, we first used backward models with a window from 0 to 15 ms to reconstruct each instrument waveform when it was attended as well as when it was ignored. The reconstruction accuracies did, however, not exhibit a statistically significant difference between the attended and the ignored case (figure 6A; *p* = 0.49 for guitar and piano).

**Figure 6.**
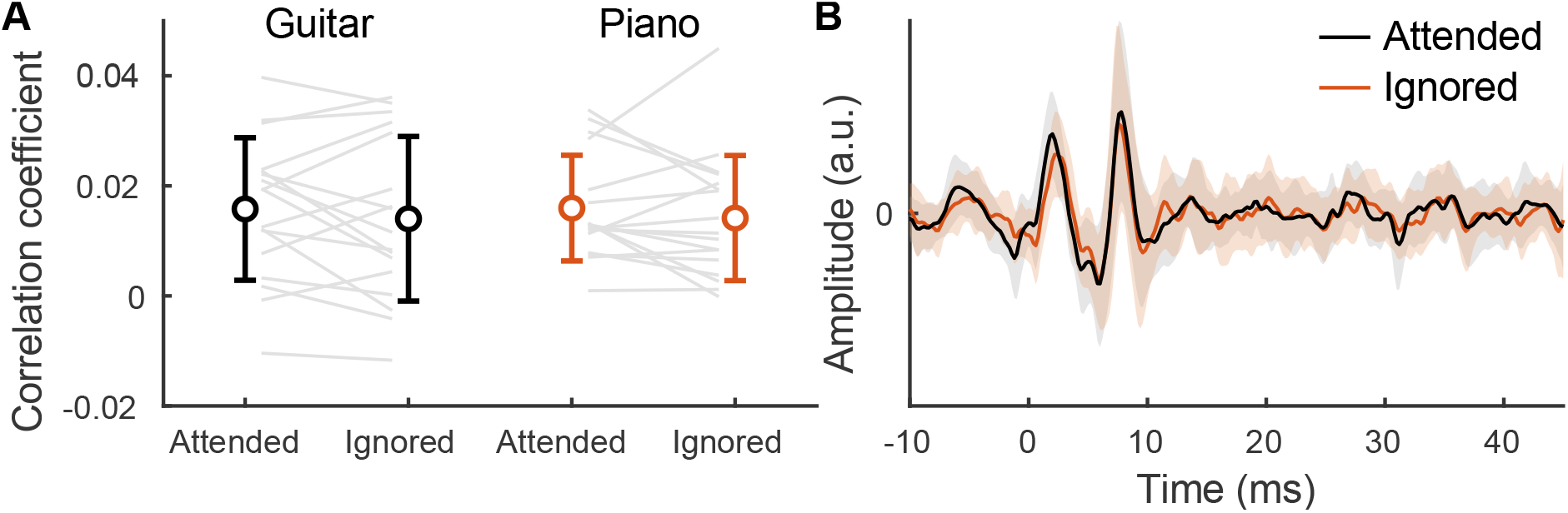
Absence of attentional modulation of neural responses. (**A**), Backward models were trained to reconstruct the stimulus waveforms for the guitar (black) and piano (red) in the CI conditions when they were attended or ignored. The reconstruction accuracies, as assessed by the correlation coefficient between the reconstructed and the original signals, did not differ significantly between the attended versus the ignored cases (*p* = 0.49 for guitar and piano). (**B**) Population-average TRFs were derived over the two CI conditions taken together for the attended (black) and ignored (red) instruments. The amplitude of the obtained TRFs did not significantly differ in the 0 ms to 15 ms region of interest. Shaded regions denote plus/minus one standard deviation across subjects around the mean TRFs.

We then computed a linear forward model that included two features, the attended and ignored instruments. The linear forward model was trained using the pooled data from the two CI conditions. The model then allowed us to compare the amplitude of the attended and ignored TRFs at each time lag from 0 to 15 ms. No significant difference between the amplitudes emerged at any temporal lag (figure 6B).

## Discussion

We showed for the first time that neural responses to the temporal fine structure of continuous musical melodies can be obtained from EEG recordings using linear convolutive models. In particular, we demonstrated that the EEG recordings could in part be predicted from the acoustic waveform (forward model, figure 3). *Vice versa*, the temporal fine structure of the musical stimuli could be decoded from the corresponding EEG recordings (backward model, figure 4). Significant responses could be obtained in most individual subjects when they were exposed to about 5 minutes of a single melodic line.

The neural response at the population level revealed further information about its origin. Indeed, the significant parts of the response, as obtained from the forward models, emerged most strongly at the latency of 7.6 ms (figure 3A). The responses at the other latencies may have reflected our use of high-pass filters for the EEG data, which spread the response in time in both directions (Widmann et al., 2015). The autocorrelation of the filtered EEG data exhibited sidelobes that are reminiscent of the structure of some of the peaks that we obtained in the neural responses (figure 2C).

The backward model showed likewise that only delays between 0 ms and 15 ms allowed for a significant reconstruction of the stimulus waveform. Together with the evidence from the forward model, these delays suggest a sub-cortical origin of the neural response, putatively in the inferior colliculus, although different sub-cortical structures may contribute as well (Bidelman, 2015, 2018; Skoe & Kraus, 2010; Sohmer et al., 1977). Recent MEG work uncovered cortical contributions to the FFR in humans (Coffey et al., 2016; Hartmann & Weisz, 2019; Ross et al., 2020), although they may be limited to frequencies below 150 Hz (Bidelman, 2018). The scalp-recorded FFR may accordingly combine multiple subcortical and cortical sources (Coffey et al., 2019). While the neural response that we have described here is arguably of subcortical origin, our use of only two EEG channels may have obstructed the observation of later cortical sources with different dipole orientations.

Neural responses can occur to both transient (e.g. clicks, onsets) and sustained (e.g. temporal fine structure) features of complex stimuli. When investigating the frequency-following response (FFR), for instance, these two aspects can be segregated by time regions (Skoe & Kraus, 2010). However, the continuous nature of the stimuli that we used here did not allow for this type of analysis. Instead, we trained a forward model with stimulus waveforms where note onsets were suppressed, and compared it to a forward model trained using the intact waveforms (figure 3A,B). The two responses were strikingly similar, suggesting that they are primarily driven by the sustained periodic oscillations of individual notes rather than their onsets. This may be expected, as these sustained oscillations constituted most of our music stimuli. In a click train, in contrast, transients dominate the temporal fine structure.

When the participants were presented with stimuli consisting of two competing instruments, they had to selectively attend to one of them, and identify vibratos that were inserted in the melodic line of that instrument. We used this task as a marker of selective attention, comparable to the use of comprehension questions in the case of speech stimuli. We found that most subjects were able to identify the target vibratos whilst ignoring the distractors (figure 5). The sensitivity index d’ was significantly larger when attending to the piano than the guitar. When attending to either instruments, the true-positive rate did not significantly differ, but the false-positive rate was lower when attending to the piano, indicating that this effect mediated the difference in d’ values. We hypothesise that since pianos cannot naturally produce vibratos, the participants may have had a bias leading to a higher propensity to attribute vibratos to the guitar. The two tasks were thus overall balanced, but attending to the piano may have been somewhat easier for the participants.

The task of attending to one of two melodic lines allowed us to investigate whether the neural response to the temporal fine structure of a particular melodic line was modulated by selective attention. Following our results on the statistical methods for obtaining this neural response, we employed backward models to reconstruct the stimulus waveform from the EEG recording, using temporal delays between 0 ms and 15 ms. We did not, however, find any significant difference between the resulting reconstruction accuracies of a melodic line when it was being attended or ignored for either instruments (figure 6A). To verify this result using a different methodological approach, we also trained a forward model that used the attended and the ignored instruments as features. Comparing the amplitude of the attended and ignored TRFs between 0 ms and 15 ms did not reveal any significant difference (figure 6B).

Our negative finding regarding attentional modulation contrasts with previous work on similar neural responses to the temporal fine structure of speech, that were found to be modulated by selective attention (Etard et al., 2019; Forte et al., 2017). It also contrasts with recent MEG work that showed that the cortical components of the FFR can be modulated by intermodal attention (Hartmann & Weisz, 2019).

These differences may point to underlying differences between music and speech. First, the two melodic lines that we used in the present work may have been difficult to selectively attend, since they originated from one musical piece, were contrapuntal, and often followed or responded to each other. The resulting interaction between the two melodic lines makes their juxtaposition rather different from that of two independent competing voices that do not interact but merely generate informational and acoustical masking. While two competing speakers may encourage selective attention and neural processing of one of them, our two melodic lines may therefore rather encourage attention, as well as neural processing, of the acoustic mixture.

Second, the subjects that participated in the competing speaker experiments effectively had a lifelong training in isolating one speaker from noise, due to the relevance of this task in daily life. As already hinted at above, we speculate that musical stimuli are instead generally perceived as a whole, and that most subjects are unfamiliar with focussing on one of several instruments. Musicians, in contrast, may in general be more familiar and trained at this task. Previous studies have indeed demonstrated that subcortical encoding of the temporal fine structure and FFR responses can exhibit long-term plasticity and that they can be modulated by musical experience (Bidelman, Gandour, et al., 2011; Bidelman, Krishnan, et al., 2011; Kraus & White-Schwoch, 2017). Similarly, musicians might exhibit attentional modulation of the neural response to the temporal fine structure of melodies, although people without musical training might not.

Finally, this study design was informed by published work analysing similar neural responses to speech (Etard et al., 2019; Forte et al., 2017; Maddox & Lee, 2018). A combination of the factors listed above may have contributed to produce neural responses differing from the ones previously reported for speech stimuli, and thus yielding no attentional modulation, or one of a much smaller magnitude. Further work is required to disentangle the potential effects of these hypotheses.

Music is a rich signal that consists of many transient and sustained features. Here, we focussed on the comparatively high-frequency neural response to the temporal fine structure. Other features, however, could be studied as well from the same stimuli, including notably cortical responses to the onsets of the notes as well as to amplitude fluctuations. Similar cortical responses to continuous speech have received significant attention in the past years, and have been shown to reflect attention (Ding & Simon, 2012a; O’Sullivan et al., 2015; Power et al., 2012) as well as semantic features (Broderick et al., 2018), surprisal (Weissbart et al., 2019) or comprehension (Etard & Reichenbach, 2019; Kösem & van Wassenhove, 2017). It has indeed been found recently that the cortical encoding of sequences of tones in a melody reflects a listener’s expectation of the upcoming notes (Di Liberto et al., 2020). Studying the interaction of such cortical responses with the subcortical activity related to the temporal fine structure that we have uncovered here may further clarify the neural mechanisms that allow us to perceive complex musical stimuli in their entirety, while also allowing us to selectively focus on a particular instrument or melodic line.

## Acknowledgement

This research was supported by the Royal British Legion Centre for Blast Injury Studies and by EPSRC grants EP/M026728/1 and EP/R032602/1. We are grateful to the Telluride Neuromorphic Cognition Engineering Workshop 2018, in particular to discussions with Carol Krumhansl, Alain de Cheveigné and Malcolm Slanely, as well as to the Imperial College High Performance Computing Service (doi: 10.14469/hpc/2232).

